# Nanoceutical Fabric Prevents COVID-19 Spread through Expelled Respiratory Droplets: A Combined Computational, Spectroscopic and Anti-microbial Study

**DOI:** 10.1101/2021.02.20.432081

**Authors:** Aniruddha Adhikari, Uttam Pal, Sayan Bayan, Susmita Mondal, Ria Ghosh, Soumendra Darbar, Tanusri Saha-Dasgupta, Samit Kumar Ray, Samir Kumar Pal

## Abstract

Centers for Disease Control and Prevention (CDC) warns the use of one-way valves or vents in free masks for potential threat of spreading COVID-19 through expelled respiratory droplets. Here, we have developed a nanoceutical cotton fabric duly sensitized with non-toxic zinc oxide nanomaterial for potential use as membrane filter in the one way valve for the ease of breathing without the threat of COVID-19 spreading. A detailed computational study revealed that zinc oxide nanoflowers (ZnO NF) with almost two-dimensional petals trap SARS-CoV-2 spike proteins, responsible to attach to ACE-2 receptors in human lung epithelial cells. The study also confirm significant denaturation of the spike proteins on the ZnO surface, revealing removal of virus upon efficient trapping. Following the computational study, we have synthesized ZnO NF on cotton matrix using hydrothermal assisted strategy. Electron microscopic, steady-state and picosecond resolved spectroscopic studies confirm attachment of ZnO NF to the cotton (i.e., cellulose) matrix at atomic level to develop the nanoceutical fabric. A detailed antimicrobial assay using *Pseudomonas aeruginosa* bacteria (model SARS-CoV-2 mimic) reveals excellent anti-microbial efficiency of the developed nanoceutical fabric. To our understanding the novel nanoceutical fabric used in one-way valve of a face mask would be the choice to assure breathing comfort along with source control of COVID-19 infection. The developed nanosensitized cloth can also be used as antibacterial/anti CoV-2 washable dress material in general.

**GRAPHICAL ABSTRACT:** 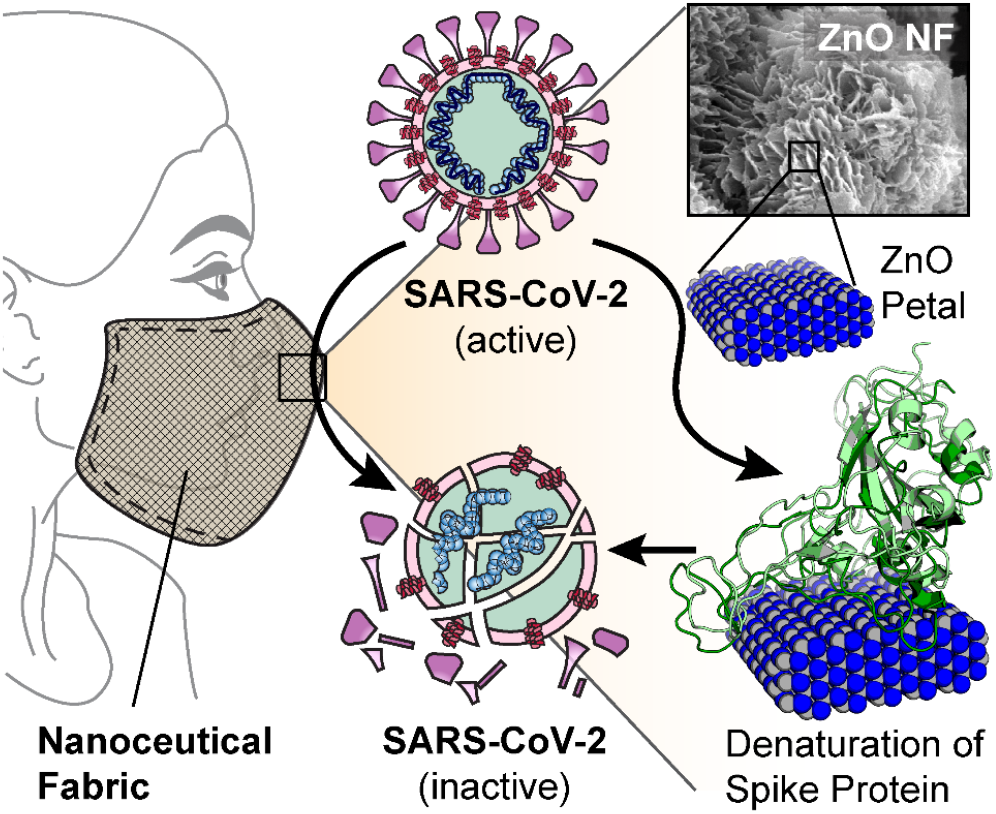

A novel nanoceutical cotton fabric duly sensitized with non-toxic zinc oxide nanoflower can potentially be used as membrane filter in the one way valve of face mask to assure breathing comfort along with source control of COVID-19 infection. The nanoceutical fabric denatures the SARS-CoV-2 spike protein and makes the microorganism ineffective.

## INTRODUCTION

The emerging coronavirus disease 2019 (COVID-19) pandemic, caused by the severe acute respiratory syndrome-related coronavirus 2 (SARS-CoV-2), has imposed a global socio-economic burden, particularly to public and private healthcare system due to unprecedented morbidity, mortality and economic disruption [1]. At the time of writing, over 81.27 million patients have been infected by SARS-CoV-2, with over 1.77 million deaths worldwide [2]. Recently, some of the vaccine candidates have received conditional approval from various authorities around the world despite numerous issues like dose, interval between two shots etc. Even after that, it will take several months to reach the common people due to challenges like regulatory issues, large-scale production and mass distribution to the public [3]. Therefore, the sole line of defence recommended by almost all the apex medical bodies throughout the world is ‘non-pharmacological interventions’ like the use of personal protective equipment (PPE) (e.g., face mask, face shield, gloves etc.), and personal hygiene (e.g., handwashing, cough and sneeze etiquette) [4–6]. As the world is ready to reopen, and most of the countries are waiting for a second or third wave of virus outbreak, the PPEs particularly face masks are becoming an integral part of day-to-day life. Although, face masks are commonly used to provide protection to the wearer (e.g., first responders), they were originally introduced to protect surrounding persons [7]. Generally, the term ‘face mask’ covers a broad range of PPEs that reduce the transmission of respiratory particles or droplets [7].

Recently, Centers for Disease Control and Prevention (CDC) warned against the use of one-way valves or vents in face masks for potential threat of spreading COVID-19 through expelled respiratory droplets [8]. However, the face masks without valves or vents can cause several health problems and severe discomfort to the wearer [9–11]. Some evidences suggest that commonly available N95 face masks can lead to changes in blood oxygen (O2) and carbon dioxide (CO2) levels when used for long periods, especially by people who are elderly, obese or have chronic obstructive pulmonary disease (COPD) [10–14]. Other adverse health effects range from increased blood pressure, increased heart rate, chest pain, and hypercapnia [15–19]. One of the feasible solution to this problem is by covering the valve or vent with a porous filter that can either trap or kill the microorganisms including viruses, alongside allow the air to flow across. Unfortunately, no study has been performed till date to design such filters or to address the problem by other means. In this regard, filters made up of natural fabrics (e.g., cotton) functionalized with antimicrobial agents could be an effective solution.

Although, a vast diversity of organic compounds with recognized antimicrobial activity have been incorporated into polymers in order to develop antimicrobial fibers, they exhibit several drawbacks related to their thermal and chemical instability, which could lead to difficulties throughout the functionalization process and also produce fibrous mats with limited antimicrobial activity due to partial decomposition of the antimicrobial agent [20]. Moreover, the massive use of antibiotics has led to another global problem, the origination of antibiotic resistance bacteria [21–24]. Thus, there is a growing need to explore novel antimicrobial agents to functionalize the natural or synthetic fabrics not only to prevent COVID-19 spread, but also for other applications like clothing materials for patients to stop hospital acquired infections, which is also a global challenge.

Consequently, nanoscale materials with unique and tuneable physico-chemical properties (i.e., higher chemical, mechanical and thermal stability) are interesting elements for preparation of hygienic surfaces. For example, silver ions and silver-based nano-compounds are highly toxic to microorganism showing strong biocidal effect various species of bacteria [25–27]. Gold nanoparticles [28, 29], bimetallic nanomaterials (e.g., Ag/Au, Ag/Pt etc.) [30, 31], graphene based materials [32] and metal oxide nanoparticles (e.g., titanium dioxide (TiO2), copper oxide (CuO), silica (SiO2) etc.) [33–35] are other examples. However, they all suffer from the problem of inherent toxicity of the nanomaterials limiting their human use [36–40]. Therefore, it is of considerable interest to design a nanoparticle functionalized fabric (i.e., nanoceutical fabric) that can efficiently kill or trap microorganisms without any harmful side effect to the user.

In this study, we have rationally designed a nanoceutical cotton fabric duly sensitized with non-toxic zinc oxide (ZnO) nanomaterial for potential use as membrane filter in the one-way valve or vent of face mask for the ease of breathing without the risk of COVID-19 spread. A comprehensive computer assisted simulation study revealed the unique potential of ZnO nanoflowers (ZnO NF) having nearly two-dimensional nanopetals in entrapment and denaturation of coronavirus spike protein (resulting into eradication of the virus), which mediates the viral pathogenesis through attachment to angiotensin converting enzyme-2 (ACE-2) receptors in human respiratory tract. Subsequently, we have synthesized ZnO NF on natural cotton fiber matrix using a one pot hydrothermal assisted approach. In depth electron-microscopic, steady-state and picosecond resolved spectroscopic studies confirm the attachment of ZnO NF to the cellulose matrix of the cotton at atomic level to develop the nanoceutical fabric filter. An exhaustive antimicrobial study using capsule containing virulent *Pseudomonas aeruginosa* as a mimic of coronavirus (as the lectins in *P. aeruginosa* shares similar homology to coronavirus spike protein) reveals excellent anti-microbial (bactericidal) efficiency of the developed nanoceutical fabric filter. To our understanding, the novel nanoceutical fabric used in one-way valve of a face mask would be the choice to assure breathing comfort along with source control of COVID-19 infection. The developed nanosensitized cloth can further be used as antibacterial (as well as anti SARS CoV-2) dress material in general to stop hospital acquired infections.

## RESULTS AND DISCUSSION

### Prediction of the potential antiviral nanostructure and model experimental organism

SARS-CoV-2 (Figure 1a) generally initiates its infection process by binding to functional receptors on the membrane of a host cell [41]. In case of humans, membrane bound ACE-2 (the functional host receptor) plays a crucial role in pathogenesis of COVID-19 providing viral entry to human cells (Figure 1b, 1c) [42, 43]. The spike (S) protein present in the outer surface of SARS-CoV-2 binds to human ACE-2 with a very strong affinity, and subsequently mediates membrane fusion leading to host cell entry of the virus (Figure 1c) [42, 44, 45]. Computed binding energy for human ACE-2 and SARS-CoV-2 S-protein from the crystal structure (PDB ID: 6lzg; Figure 1d) of the complex was found to be −20.28 kcal mol^−1^. Following minimization of the complex under OPLS (Optimized Potentials for Liquid Simulations) [46] force field, the binding energy drops to −29.83 kcal mol^−1^. This strong binding energy can be attributed to the large surface area at the protein-protein interface and several favourable intermolecular interactions including hydrogen bonding, ionic and hydrophobic interactions. Hence, it is difficult to develop small molecule competitive inhibitors that can disrupt such an elaborate molecular recognition. Therefore, we looked into the prospective nanomaterials having similar magnitude as ACE-2 protein along with the ability to compete for the viral S-protein binding. In this regard, our primary choice was ZnO considering its biocompatibility and known antimicrobial and antiviral effect [47–53]. In a computer simulation, spherical ZnO nanoparticle (~30 nm diameter, similar dimension of ACE-2) was constructed and its interaction with the viral S-protein was studied (Figure 1e). The spherical nanoparticle provided a convex surface for interaction similar to the ACE-2 receptor protein. Since, the geometry of the nanosurface at the binding region is known to affect the protein-nanomaterial interaction [54, 55], we further simulated the interaction of S-protein with the flat sheet-like ZnO surface having various exposed crystal facets (<100>, <101> and <002>) as observed in X-Ray diffraction (XRD) pattern (*vide infra*).

**Figure 1.**
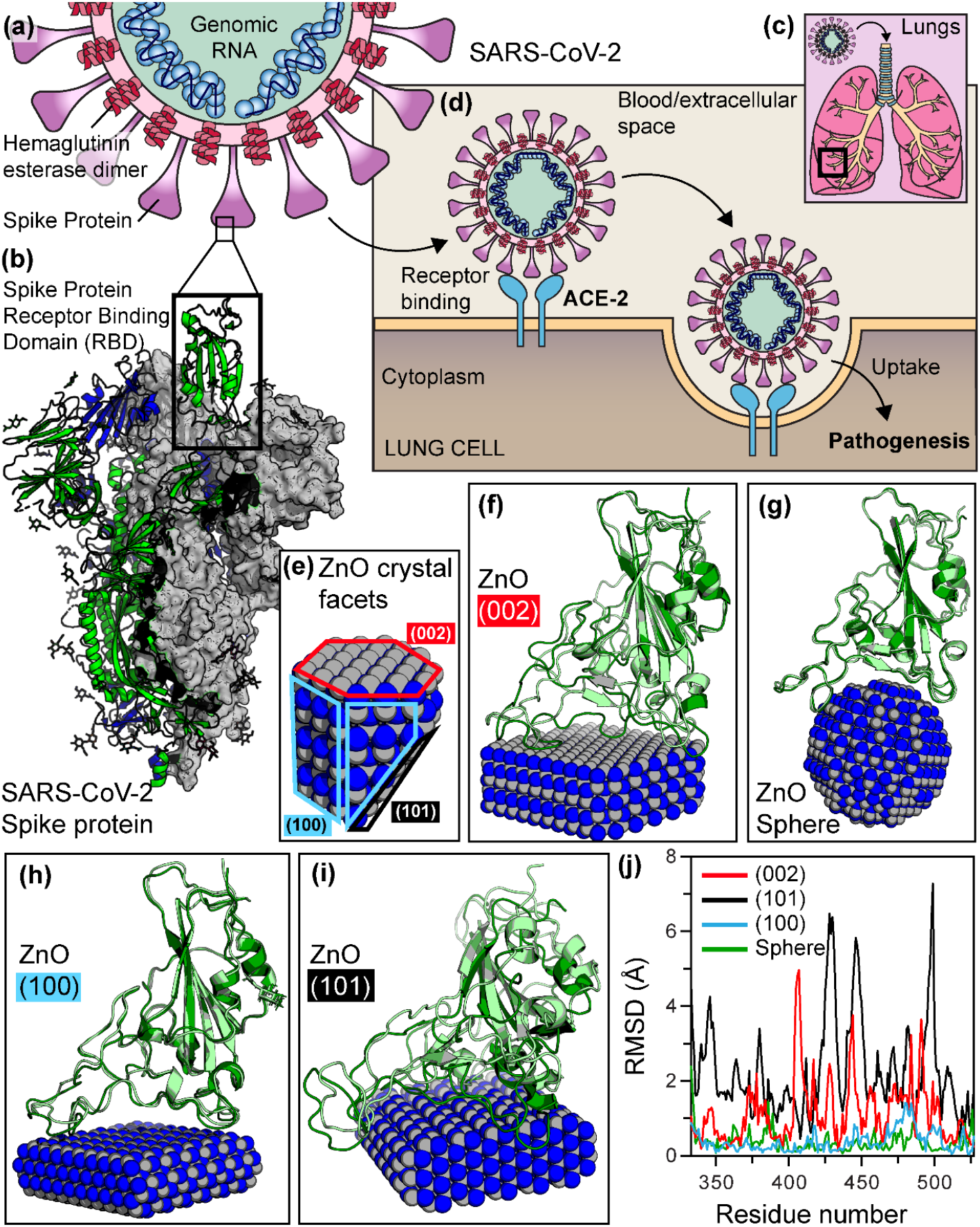
Interaction of SARS-CoV-2 spike protein receptor binding domain (RBD) with ZnO nanofacets leads to denaturation of spike RBD. (a) Schematic representation of SARS-CoV-2 structure. This is an enveloped, positive sense RNA virus having spike proteins at the outer surface. (b) Side view of the pre-fusion structure of the SARS-CoV-2 spike protein with a single RBD in the up conformation. (c) SARS-CoV-2 infection through human respiratory tract. (d) Internalization of SARS-CoV-2 by membrane fusion. The spike protein binds to the angiotensin-converting enzyme 2 (ACE-2) of the host cell via RBD and further releases its RNA to exert pathogenesis. (e) Crystal facets of Wurtzite zinc-oxide (ZnO) nanostructure used in computational studies. (f - i) Binding of spike RBD (molecular docking) with different facets of ZnO nanostructure as well as spherical ZnO nanoparticle. The spike protein strongly binds to the ZnO crystal facets and denatures upon interaction. Native structure of the protein is represented by light color, while dark colors represents the denatured structure after interaction. (j) Root mean square deviation (RMSD) values of each amino acids after energy minimization under optimized potentials for liquid simulations (OPLS) shows the degree of denaturation.

As evident from molecular docking study, the SARS-CoV-2 S-protein can bind favorably with both the ZnO nanosphere as well as on the flat facets (<100>, <101> and <002>) of ZnO nanostructure with binding energies: −17, −11, −21 and −16 kcal mol^−1^, respectively. Binding on <101> ZnO nanosurface was very much comparable with that of ACE-2. After the energy minimization of the docked complexes, the binding energies were found to be −21, −13, −40 and −22 kcal mol^−1^, respectively. Figure 1e-1i shows the bound conformations of spike protein with the ZnO nanosphere and various facets on ZnO nanosurfaces. Interestingly, when structural deviations (in terms of root mean square deviation, RMSD) were studied and compared between the docked and the energy minimized conformations of the spike RBD (Figure 1j), large structural perturbations become evident in the binding regions of SARS-CoV-2 S-protein due to binding on ZnO nanofacets. Least change was observed in case of nanosphere, whereas most change occurred on <101> facet of ZnO. Since, structure and function of proteins are highly correlated, infliction of a large structural perturbation in protein structure by an inhibitor, or ligand is generally associated with its loss of function [46]. The observed *in silico* structural changes, therefore, indicate that the viral S-protein may lose its pathological function if it comes in contact with ZnO nanosurface.

Bearing in mind the transmissibility and pathogenicity of SARS-CoV-2 as well as unavailability of the SARS-CoV-2 strain and biosafety requirements for experimental studies, we looked for similar organisms as an alternative model preferably of bacterial origin having adhesion proteins analogous to viral S-protein. Lectin-A (LecA) protein of *P. aeruginosa* (Figure 2a) was found to show high degree of similarity in the binding site residues as evident from the amino acid sequence alignment depicted in Figure 2a-2e. In the largest 32 amino acid overlap region (14-45:149-180), the two proteins showed 28.1% identity, and 62.5% similarity with Waterman-Eggert score of 52. Detailed sequence alignments of the Spike RBD and LecA protein are given in supporting information. Apart from the structural similarity, LecA resembles the function of viral S-protein, and mediates bacterium-host cell recognition and adhesion which are the critical steps in initiating *P. aeruginosa* pathogenesis [56]. Interestingly for both SARS-CoV-2 and *P. aeruginosa*, lungs are the primary target organs, and both of them cause similar acute and chronic lung infection [56–58]. Therefore, considering the structural and functional resemblances in a possible example of molecular mimicry, *P. aeruginosa* LecA was chosen as an alternative to SARS-CoV-2 S-protein for subsequent computational and experimental binding studies with the ZnO nano structures.

**Figure 2.**
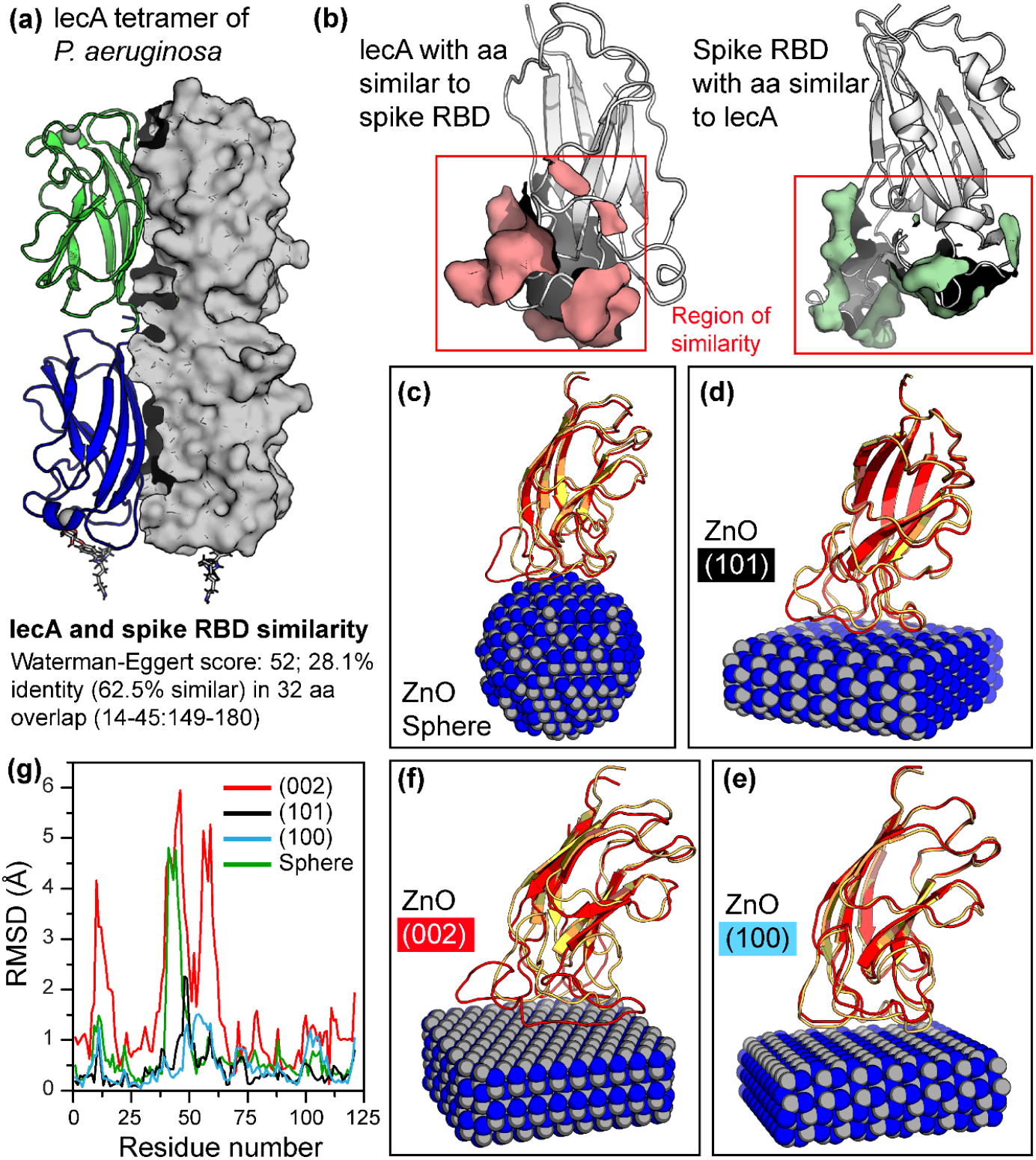
Interaction of a mimic of SARS-CoV-2 spike RBD, lecA of *P. aeruginosa,* with ZnO nanofacets leads to denaturation. (a) Tetrameric crystal structure of *P. aeruginosa* lecA. (b) Computed structural similarity between lecA and spike RBD. (c - f) Binding of lecA (molecular docking) with spherical ZnO nanoparticle as well as different facets of ZnO nanostructure. LecA strongly binds to the ZnO crystal facets and denatures upon interaction. Native structure of the protein is represented by light color, while dark colors represents the denatured structure after interaction. (j) RMSD values of each amino acids after energy minimization under OPLS shows the degree of denaturation.

Molecular docking study illustrates that the P. aeruginosa LecA protein can bind favorably with the ZnO <100>, <101> and <002> nanofacets as well as on the ZnO nanosphere with comparable binding energies: −9, −11, −12 and −11 kcal mol^−1^, respectively. After the energy minimization of the docked complexes, the binding energies were found to be −11, −14, −26, and −17 kcal mol^−1^, respectively. Figure 2c-2f shows the bound conformations of LecA protein with the ZnO nanosphere and nanofacets, respectively. Structural deviations in terms of RMSDs were studied and compared between the docked and the minimized complexes and graphically represented in Figure 2g. It is evident from Figure 2g that large structural perturbation, similar to the viral S-protein, takes place in the binding regions of LecA protein on the ZnO nanofacets.

Thus, the computational binding study shows that ZnO nano surface can trap the adhesion proteins of SARS-Cov-2 as well as that of the *P. aeruginosa*. The sheet-like flat ZnO nanostructures not only binds these adhesion proteins (i.e., viral S-protein and LecA), but also impart large scale structural perturbation in the bound protein (i.e., denaturation) leading to functional impairment and disinfectant effect. Therefore, based on the computational findings *P. aeruginosa* (having LecA) was chosen as model SARS-CoV-2 mimic, and ZnO sheet-like nanostructure was selected as anti-microbial agent for subsequent experimental validation.

### Designing of natural cotton fabric functionalized with ZnO nanoflower featuring flat sheet-like petals

Encouraged by the *in silico* studies, we intended to decorate the natural cotton fabric with flat ZnO sheet like structures, which eventually self-assembled into more complex rose-like ZnO nanoflower during hydrothermal-assisted low-temperature one pot synthesis process (detailed in ‘Methods’ section). The scanning electron micrographs (SEM) confirm the formation of flower like morphology of ZnO on the surface of the cotton fiber (Figure 3a-3d). Figure 3e-3g shows the SEM of the bare cloth. The flat sheet like petals of the flower have a length of ~600-850 nm, and a diameter of ~150-300 nm with an aspect ratio of ~3 (schematically represented in Figure 3h). It should be noted that the surface of the petals is not smooth, rather coated with small granules of nanoparticles. The EDAX analysis confirms presence of ZnO in the nanostructure (data not shown). TG-DTA analysis shows ~9% (w/w) loading of ZnO NF to cellulose fibers (Supplementary Figure S1).

**Figure 3.**
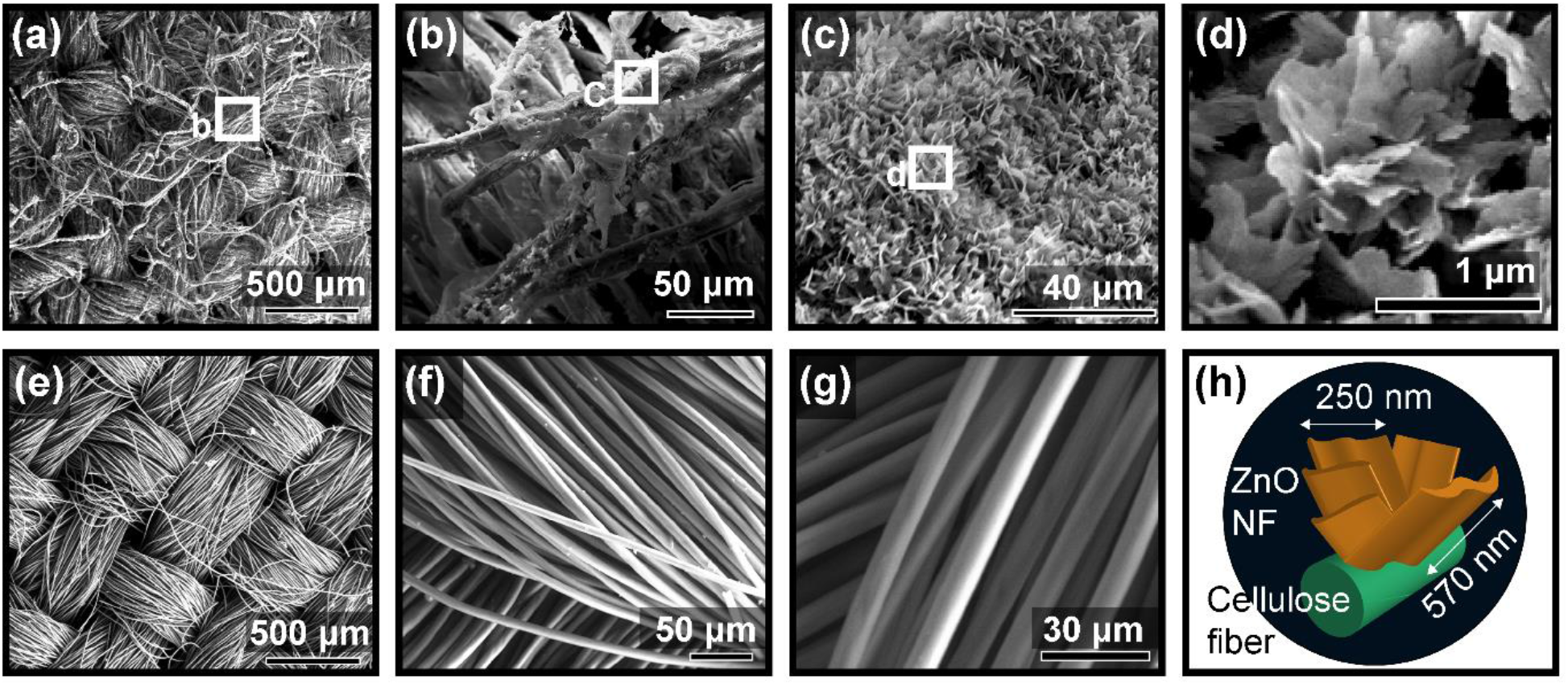
Scanning electron micrographs (SEM) of the developed ZnO nanoflower (ZnO NF) decorated cotton cloth at different magnification. (a-c) Low magnification SEM of ZnO NF decorated cotton cloth. (d) High magnification SEM shows the self-assembled petal like structures of a single nanoflower (e-g) SEM of bare cotton cloth showing the constituent cellulose fibers. (h) Schematic representation of the ZnO NF as observed under SEM

At the outset, we investigated the growth of ZnO phase on the cotton clothes using X-ray diffraction (XRD) technique. With 5 hrs of intermediate growth time, the ZnO phase is not distinguishable properly as the prominent XRD peaks are suppressed by the broad hump of cellulose within the Bragg angle (2*θ*) range between 31° and 38° in Figure 4a. Nevertheless, the signature of (1 0 1) plane of ZnO at ~36.2° can be traced from the XRD pattern. The growth of ZnO becomes quite dominant when the growth time is increased to 15 hrs. In this case, the diffraction peaks appearing at 2*θ* of 31.8°, 34.4°, 36.1°, 47.4°, 56.6°, 63.0°, 68.0°, and 69.2° represent the (1 0 0), (0 0 2), (1 0 1), (1 0 2), (1 1 0), (1 0 3), (1 1 2), and (2 0 1) planes of ZnO nanocrystals, respectively. The observed peaks are in good agreement with those for hexagonal closed-packed ZnO having wurtzite structure (lattice constants *a*=3.249, *c*=5.206 Å; space group P63mc (186)) as reported earlier (JCPDS 36-1451) [59]. Sharp diffraction peaks indicate good crystalline quality of the synthesized nanoflowers, and the absence of any additional peak directs towards the high purity of the material. The highly intensified peak at 36.1° indicates the dominance of (1 0 1) plane in the material.

**Figure 4.**
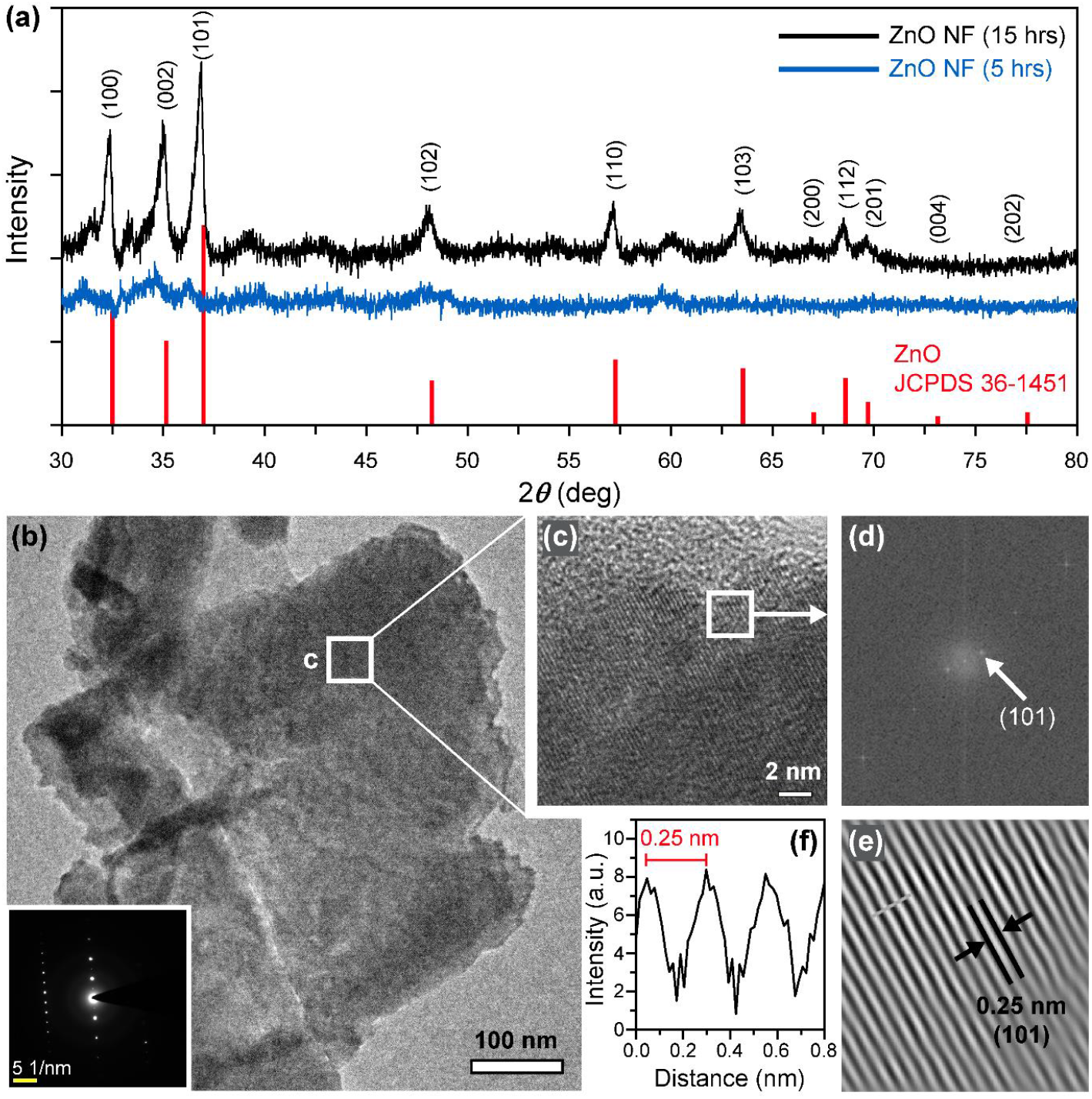
Characterization of the ZnO NF decorated cotton cloth. (a) X-Ray diffraction (XRD) pattern of the ZnO NF decorated cotton cloth at different synthesis phase. (b) Transmission electron micrograph (TEM) of a single fully grown petal. Inset shows a selected area electron diffraction (SAED) pattern of the ZnO petal. (c) HRTEM image of a part of the petal indicated as white box in (b). (d) Corresponding fast Fourier transform (FFT) image. (101) plane is indicated. (e) Reconstructed lattice image from the FFT. (f) Inter-planar spacing plot shows d-spacing of ~0.25 nm.

The TEM image (Figure 4b) shows the size of a single ZnO NF petal to be ~600 × 300 nm. The HRTEM image taken at the centre of the petal indicates the presence of (101) plane (Figure 4c). The presence of distinct bright spots in the SAED pattern (Figure 4d) corresponds to the crystalline nature of the petals. Fast Furier transformation (FFT) shows that the crystallite is dominated by (1 0 1) plane of ZnO, with an lattice fringe spacing of 0.25 nm (Figure 4e & 4f). Further, as shown in Figure 5a, the formation of nanorod like morphology can be observed when the reaction time is reduced (i.e., 5 hrs.). The surface of the nanorods appears to be rough and can be originated from the presence of large number of crystallites. The SAED pattern shows small spots within diffused ring and reveals the polycrystallinity of the nanorods as each spot arises from the Bragg reflection from individual crystallites (Figure 5a-inset). The rings can be indexed as (101), (102) and (200) of wurtzite phase of ZnO, which exactly correlates with the XRD pattern (Figure 4a). Next the close look of the nanorod at different locations (marked as c, d in Figure 5b) reveal the presence of (101) and (002) planes of ZnO with interplanar spacing ~ 0.25 and 0.26 nm respectively (Figure 5c-5g). The exhibition of polycrystalline nature of the nanorods with various crystallites is different from nanorods grown on crystalline surfaces (e.g. Si). Thus it can be anticipated that the use of cotton substrate in the synthesis process play a big role in forming nanorods with exposed crystallites on the surface (Figure 6). In fact, the functional groups present in the cellulose fibres act as nucleation sites for the growth of nanorods (Step 1 in Figure 5). It can be noted that in HMT based hydrothermal synthesis, Zn(OH)_2_ acts as the nucleation agent according to the following equations [60]:

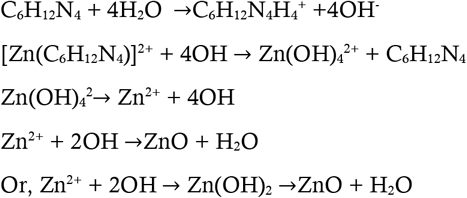

**Figure 5.**
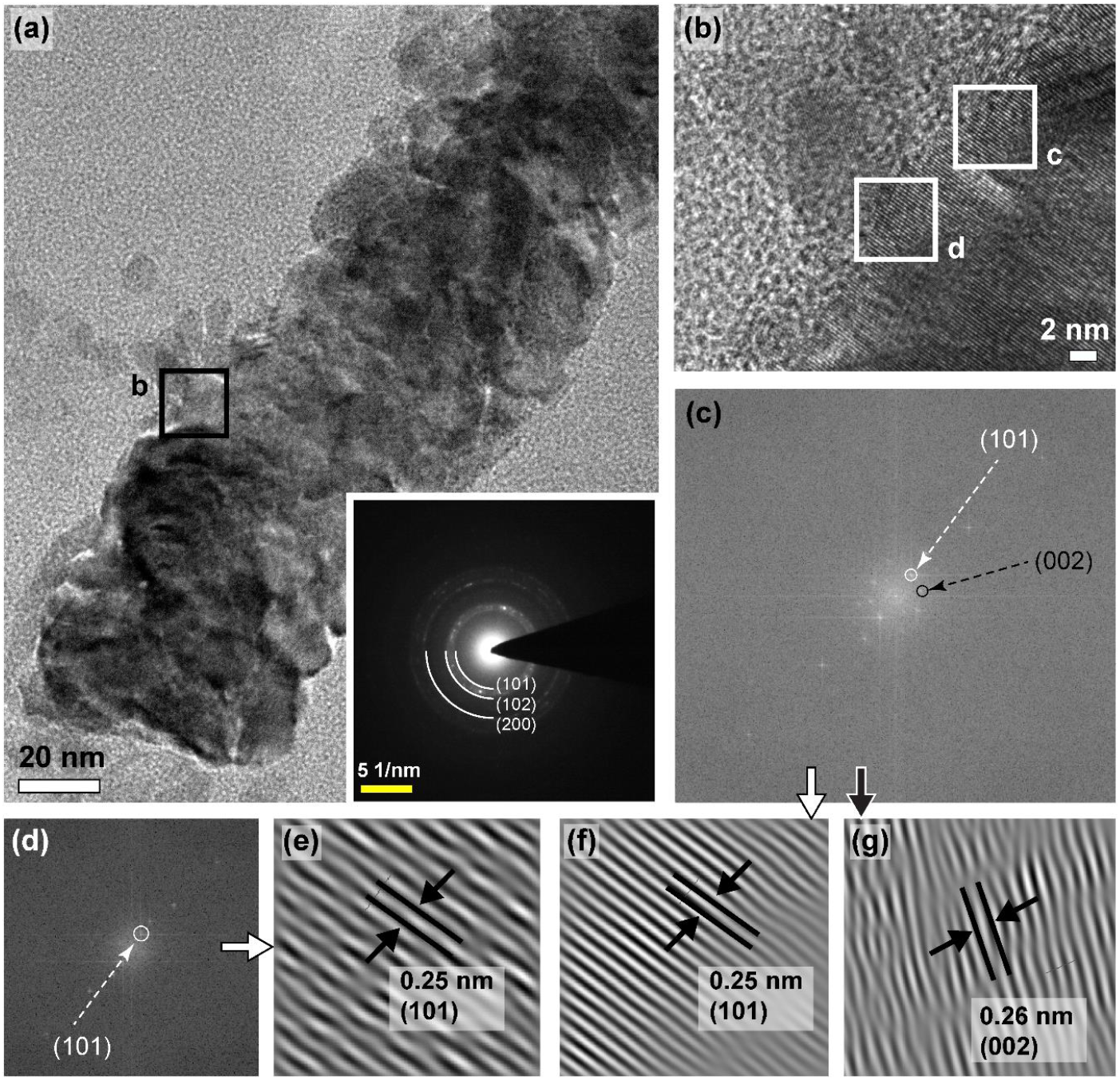
TEM image of ZnO NF petal at intermediate growth time (5 hrs). (a) TEM of a single growing petal. Inset shows the SAED pattern of the growing petal. (b) HRTEM of the petal. (c - d) Corresponding fast Fourier transform (FFT) image from different areas indicated as white box in (b). (101), and (002) plane is indicated. (e - g) Reconstructed lattice image from the FFT shows the fringe pattern and d-spacing.

**Figure 6.**
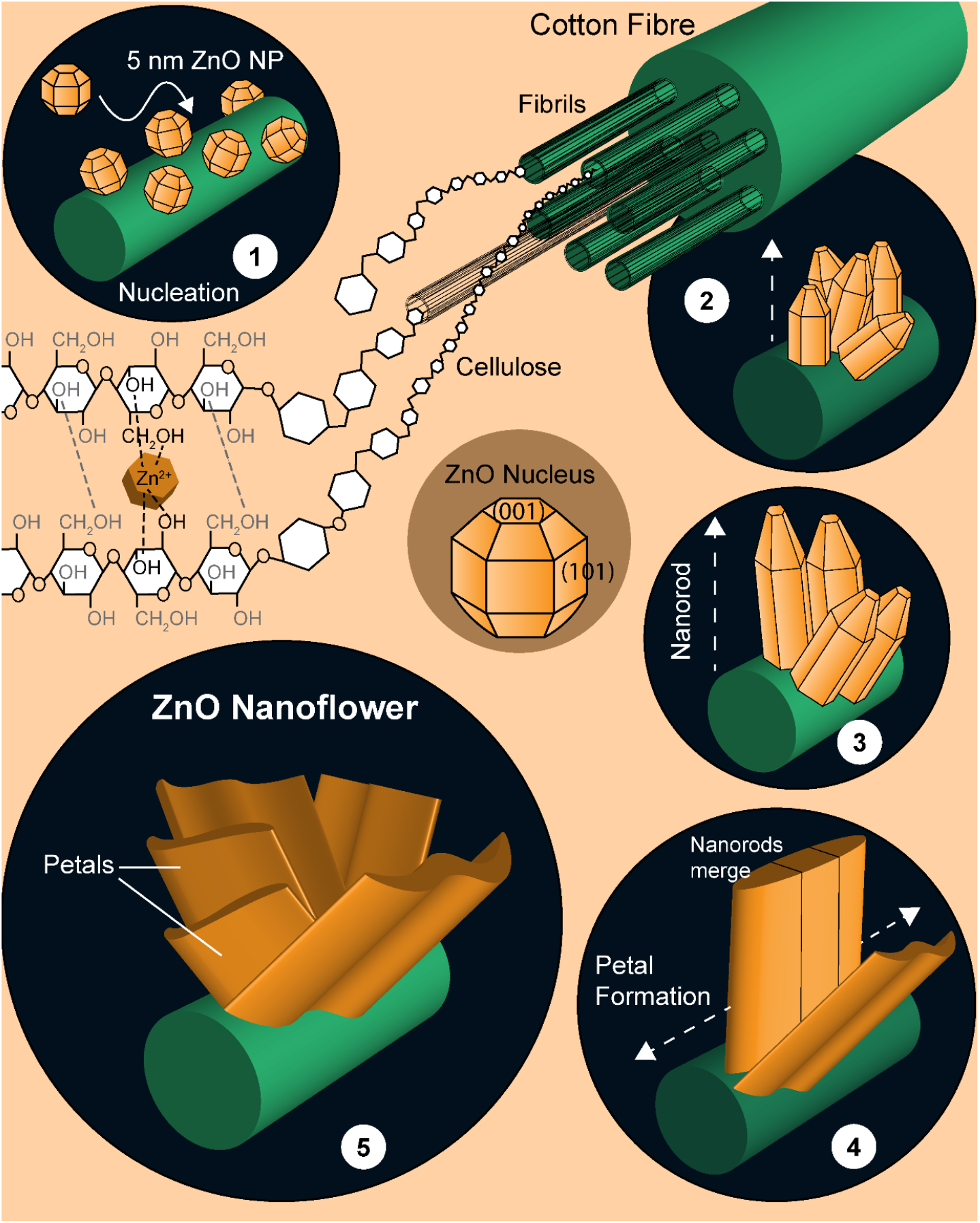
Schematic representation of the different phases of ZnO nanoflower growth on cotton cloth constituted with cellulose fibers.

In the present investigation, the abundant and random distribution of these functional groups based nucleation sites on the cellulose fibre surface provides a continuous platform for ZnO growth along the direction of fibres (Step 1 in Figure 6). As the nucleation and subsequent growth process occurs on the outer circumference of the cellulose fibres, the fibre plays the role of template to form nanorod like one dimensional structure (Step 2 in Figure 6). Now the random distribution of the functional groups leads to the randomly oriented nucleation sites and consequently the growth is evident to be different in different directions. The observation of the rough surface with different crystallites of (101) and (002) planes, is the consequence of such growth process.

Now it is well established that in ZnO crystals the surface energy (per mole) varies with direction as γ_[0001]_< γ_[11.20]_ γ Y_[10-10]_ [61]. Hence, the growth of (002) plane along [0001] direction is highly favourable from thermodynamic point of view. Thus, the nanorods with exposed (002) planes offer chances for further growth. As a result, with increased reaction time, the nanorods suffer two dimensional growth i.e. along horizontal as well as vertical directions to the surface of the fibre leading to the development of petal like feature (Step 4 in Figure 6). To ensure this fact, TEM analysis has been performed on four different edges of the fully grown petal. In none of these areas, any signature of (002) plane has been found. It is well-known that a plane with higher growth rate disappears quickly [62]. Thus the disappearance of (002) plane confirms the speculated lateral growth of the nanorods to form the petals. Subsequent self-assembly of such petal like features leads to the formation of flower like morphology of ZnO at higher reaction time (Step 5 in Figure 6).

Figure 7a shows the absorbance spectrum recorded from bare cotton cloths and ZnO NF decorated cotton cloths. As the constituent of cotton fiber is cellulose, the observed absorption bands at 200–220 and 270–290 nm could be assigned to oxidized xylans and monocarboxyl celluloses due to presence of carboxyl and carbonyl groups, respectively [63]. Furthermore, the absorption bands at 230–250 and 290–320 nm could be assigned to heteroaromatics of the furan and pyron type, and the low intensity absorption shoulder above 300 nm to conjugated heteroaromatics [63]. Functionalization with ZnO NF lead to the appearance of characteristic ZnO absorption edge at 365 nm which is blue shifted by 8 nm relative to the bulk exciton absorption (373 nm) [64]. Correspondingly, the ZnO NF exhibited a slightly higher optical energy gap (E_g_) of 3.44 eV with respect to bulk ZnO (3.37 eV) (Figure 7a-inset). The slight change in the absorption edge and Eg can be well explained by the oriented attachment of multiple ZnO nanorods in flower like morphology [64, 65].

**Figure 7.**
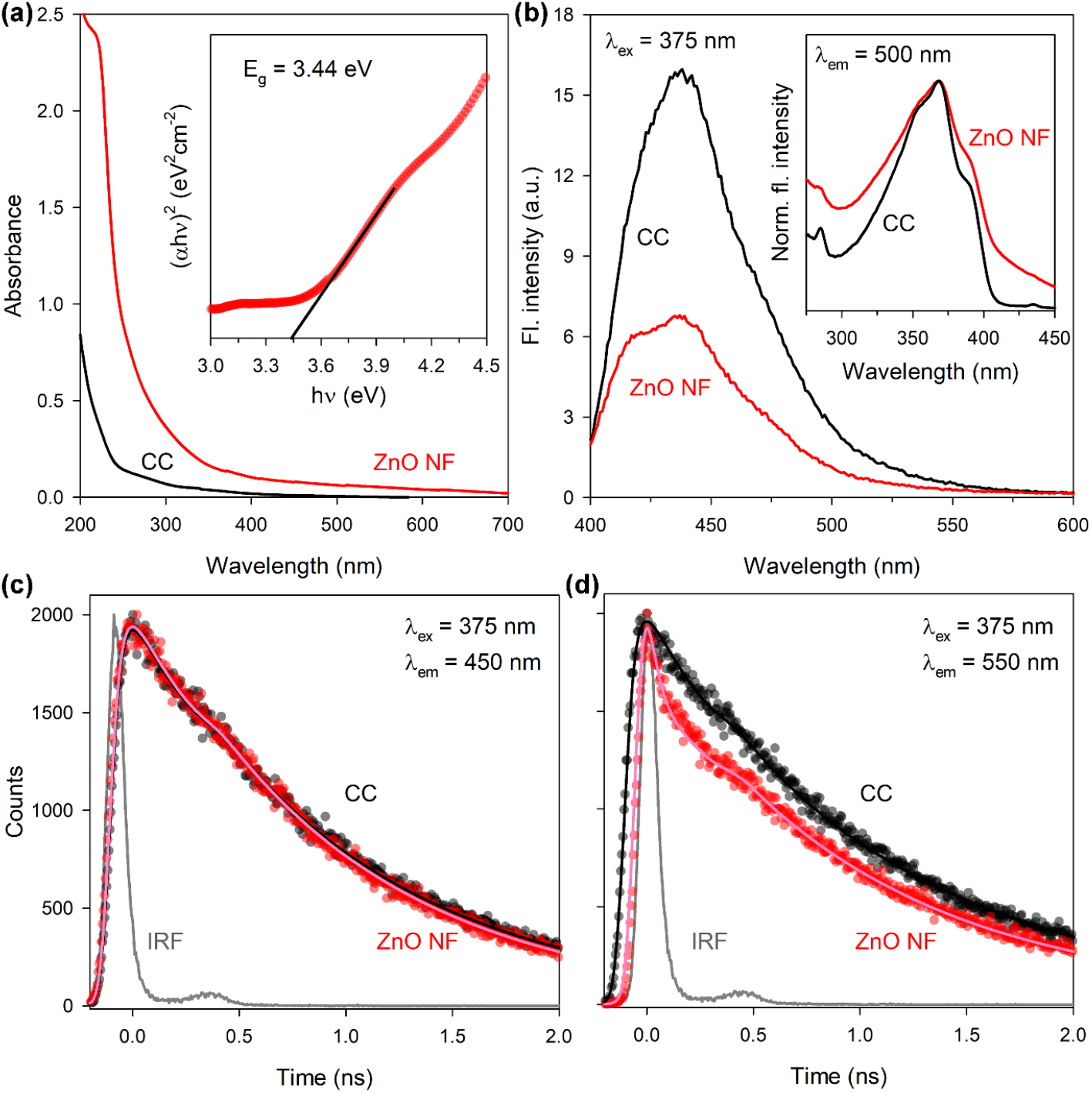
Spectroscopic characterization of the ZnO NF decorated cotton cloth. (a) UV-Vis absorbance spectra of ZnO NF decorated cotton cloth and bare cotton cloth. Inset shows Tauc’s plots for determination of the band gap energy. (b) Steady state fluorescence emission spectra shows quenching of cotton cloth fluorescence (originated from constituent cellulose) upon ZnO NF decoration. Inset shows corresponding change in fluorescence excitation spectra. (c) & (d) Picosecond resolved transients show quenching of fluorescence lifetime of the constituent cellulose by ZnO NF at longer wavelength.

The fluorescence emission spectra of cotton cloth in the solid state (Figure 7b) shows an intense peak around 430 nm and a broad shoulder ranging from 470-560 nm when excited at 375 nm. The observed emission pattern of cotton cloth exactly matches to the previously reported photoluminescence characteristics of cellulose fibers [63, 66]. Gavrilov and Ermolenko (1966) [67] who studied photoluminescence of cellulose in great details, have suggested that the broad emission band of cellulose originates from three different types of excitation centers. Presence of multiple excitation maxima (i.e., 265 nm, 350 nm, and 390 nm) in the excitation spectra of bare cotton cloth (Figure 7b-inset) further supports their findings. Functionalization with ZnO NF significantly quenched the emission of cellulose constituent (Figure 7b), which indicates possible interaction between ZnO NF and cellulose fibers. The observed increase in the λ_em_ 410 nm/λ_em_ 450 nm ratio could be due to a possible interaction between Zn-vacancy state of ZnO NF and the OH groups of cellulose fibers (O=C–O–Zn), as previous studies suggests that the presence of carboxylic groups in the anhydroglucose units shifts the emission maxima of cellulose to shorter wavelength (blue-shift) [63, 66, 68]. A detailed computational study (i.e., molecular docking) further supports the strong binding of ZnO NF with the cellulose fiber, mostly through hydrogen bonding. The calculated binding energies per unit area of cellulose fiber with different facets of ZnO nanostructure were found to be −21 kcal mol^−1^ nm^−2^ (sphere), −43 kcal mol^−1^ nm^−2^ ((002) facet), −41 kcal mol^−1^ nm^−2^ ((100) and (110) facet), and −39 kcal mol^−1^ nm^−2^ ((101) facet) (Supplementary Figure S2). The highly negative binding energies indicate stronger attachment between the cellulose fiber and ZnO NF.

To get further insight of the quenching mechanism, we studied excited state fluorescence lifetime of cellulose (i.e. cotton cloth) and ZnO NF decorated cellulose. Interestingly, the fluorescence lifetime of both the samples remained unchanged at 450 nm (λ_ex_ 375 nm) (Figure 7c), while a decrease was observed at 550 nm (λ_ex_ 375 nm) upon ZnO NF decoration (Figure 7d). This could well be interpreted by the existence of several luminescent centers (originated from the presence of conjugated chromophores of various origins) in the cellulose [67]. One possible mechanism behind the observed decrease in the fluorescence lifetime of ZnO NF decorated cellulose could be due to the photoinduced electron transfer (PET) from the LUMO level of the low energy excitation centers (near red end) of cellulose to the conduction band of ZnO, resulting in the luminescence quenching of cellulose [69]. Presence of a very fast component (i.e., 25 ps) in the excited state lifetime decay of ZnO NF decorated cellulose further supports our conjecture. In contrast, due to the mismatch in the energy levels, the high energy excitation centers (near blue end) of cellulose cannot undergo such electron or energy transfers. The quenching in this region is static in nature and may be originated due to ground state interaction between ZnO NF and cellulose which we explained in the earlier section. In summary, the results of our optical studies confirms the attachment of ZnO NF to the cellulose fibers of the cotton cloth.

### Antimicrobial activity of ZnO NF functionalized cotton fabric

The antimicrobial activity of the ZnO NF decorated cotton fabric was evaluated against the SARS-CoV-2 mimic pathogenic capsule containing (having LecA, the mimic of viral spike protein; *vide* computational studies in the earlier section) gram negative model bacteria *P. aeruginosa.* To directly assess the antimicrobial effect, after 1 hr of incubation (in dark, 37°C) over the ZnO NF decorated cotton cloth and bare cotton cloth, the *P. aeruginosa* was transferred and cultured on a Luria–Bertani (LB) agar media (i.e., the plate count assay) for 12 hrs (at 37°C). The original bacterial inoculum, which was employed over the fabrics for initial incubation served as control. As shown in the photographs of the agar plates after 12 hrs (Figure 8a), a clear killing activity was observed in case of ZnO NF functionalized cotton fabric resulting into very few colonies. Colony count shows (Figure 8b) that in ZnO NF functionalized cotton fabric the bacterial growth was ~79% lower compared to control (*P*<0.0001; F (2,15)=303.2, one-way ANOVA) and ~54% lower compared to bare cotton fabric (*P*<0.0001; F (2,15)=303.2, one-way ANOVA).

**Figure 8.**
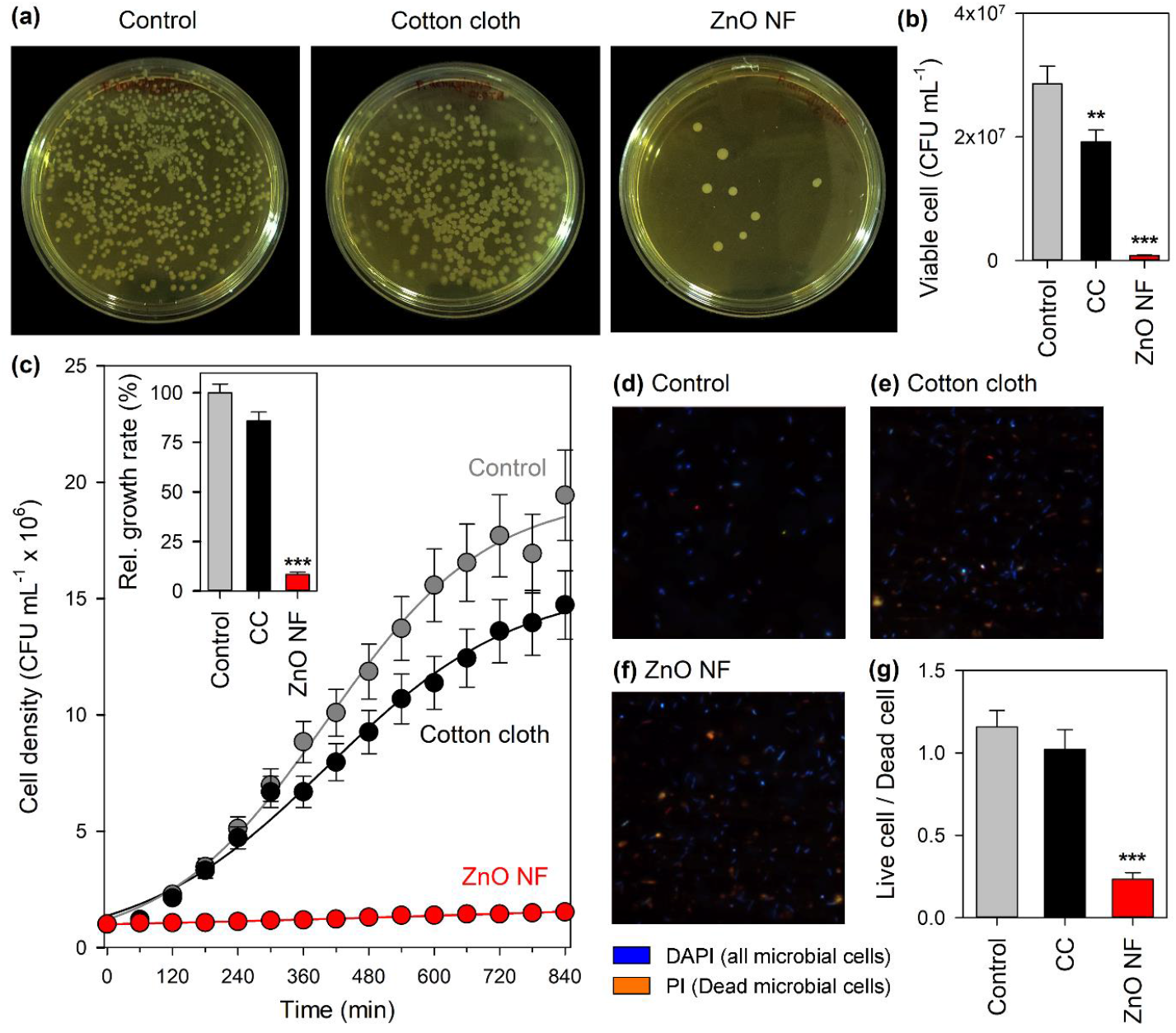
Antimicrobial activity of ZnO NF decorated cotton fabric on *P. aeruginosa,* a mimic of SARS-CoV-2. (a) Digital photographs of P. aeruginosa colonies on agar plate after treatment with ZnO NF decorated cotton cloth and bare cotton cloth. (b) Bacterial cell count after 12 hrs. of incubation. (c) Growth pattern of P. aeruginosa treated with ZnO NF decorated and bare cotton cloth. Inset shows relative growth rates. (d - f) Micrographs of bacteria after DAPI-PI differential staining. (g) Corresponding ratio of live vs dead cells.

Further, the antibacterial effect of ZnO NF decorated cotton cloth on the *P. aeruginosa* growth kinetics was evaluated in liquid LB media. The bacterial growth was monitored by the measurement of the optical density at 600 nm (OD_600_), as a reflection of the bacterial concentration in the culture. After 1 hr of incubation over respective fabrics (inoculation at 1 × 10^6^ CFU/mL), we started monitoring the growth kinetics. The original bacterial inoculum, which was employed over the fabrics for initial incubation served as control. Figure 8c, clearly indicates that ZnO NF decorated cotton cloth caused complete inhibition of growth and this was even continued for 14 hrs. The bare cotton cloth also showed a weaker growth inhibition. The growth rate was ~10 times lower in case of ZnO NF functionalized cotton fabric compared to both control (*P*<0.0001; F (2,15)=1062, one-way ANOVA) and bare cotton fabric (*P*<0.0001; F (2,15)=1062, one-way ANOVA) (Figure 8c-inset).

Our computational studies have suggested that ZnO NF can simultaneously bind and denature *P. aeruginosa* LecA (SARS-CoV-2 spike protein mimic), resulting into disruption of bacterial cell membrane, which eventually is responsible for the observed antimicrobial assay. In order to confirm that ZnO NF disrupt the bacterial membrane, *P. aeruginosa* cells were treated with PI after incubation with ZnO NF decorated cotton fabric. In is worth mentioning here that PI is a small-molecule dye that upon binding to double-stranded DNA fluoresces red upon excitation, and it cannot cross an intact cytoplasmic membrane [70, 71]. We expected that ZnO NF-induced membrane disruption would lead to an increase in nucleic acid staining by PI compared to that of control or bare cotton cloth treated ones. As shown in Figure 8d & 8e, control bacteria, as well as bacteria incubated with bare cotton fabric remained unstained to PI. In comparison, intracellular PI staining was clearly visible in cultures exposed to ZnO NF decorated cotton fabric (Figure 8f). We further evaluated the number of viable cells using differential staining with DAPI and PI. As discusses earlier, the PI generally stains the membrane disrupted cells whereas, DAPI stains all nuclear material irrespective of their viability. The differentially stained images of control and cotton fabric inoculated bacteria shows more number of viable cells than dead cells (ratio ~ 1.2) (Figure 8g). In contrast, the number of living cells were significantly reduced in case of ZnO NF functionalized cotton fabric (~0.45) (Figure 8g).

Thus, our antimicrobial studies experimentally provide evidence that ZnO NF can induce membrane damage, resulting into a loss of membrane potential leading to inhibition of PI staining. It further depicts that ZnO NF is an effective anti-microbial agent particularly against SARS-CoV-2 mimicking microorganisms.

### Washability and reusability of the ZnO NF functionalized cotton fabric

For regular use as antimicrobial hospital dress material (or, PPE), washability and reusability of the fabric are the two major factors. To address this issue, we have washed the fabric material with a mild detergent, as well as with a harsh detergent in tap water. Even after 50 washes with mild, and 10 washes with harsh detergent, the ZnO NF coating of the cotton fibers remained almost intact as observed under SEM (Supplementary Figure S3). The antibacterial activity was also sustained in both the cases as compared to the original ZnO NF functionalized fabric (Supplementary Figure S4). Therefore, the ZnO NF functionalized cotton fabric was found to be highly reusable with sufficient washability.

## CONCLUSION

In conclusion, we successfully functionalized the commonly available cotton fabrics with ZnO nanoflowers (ZnO NF). The ZnO NF functionalized cotton fabrics showed significant antimicrobial activities against SARS-Cov-2 mimic model pathogenic microorganism as depicted in our detailed antimicrobial assays. The ZnO NFs were able to destroy both microbial membrane leading to inhibition of infection. A detailed computational study along with *in vitro* studies using viral spike protein mimicking protein and bacteria showed the trapping ability of the fabric. As a proof of concept, we designed a laboratory grade prototype respirator for using in common N95 masks using the ZnO NF functionalized cotton fabrics. The porous respirator helped to solve the problem of CO_2_ rebreathing and prevented spread of microbes through the pores. The ZnO NF functionalized cotton fabrics can further be used to design comfortable, washable anti-microbial PPE, which is an urgent need of today.

## MATERIALS AND METHODS

### Synthesis of ZnO NF functionalized cotton cloth

Zinc nitrate hexahydrate (Zn(NO_3_)_2_, 6H_2_O; Sigma-Aldrich), hexamethylenetetramine (C_6_H_12_N_4_; Aldrich) were used as the starting materials for a low temperature hydrothermal synthesis of ZnO NFs on cotton fiber matrix.

For synthesis of ZnO functionalized cotton cloth, at first, seed layer ZnO nanoparticles of ~5 nm in size were prepared in ethanol. In a typical synthesis process, to a 2 mM zinc acetate (Zn(CH_3_COO)_2_·2H_2_O; 20 mL in ethanol) solution heated at 55 °C, 4 mM sodium hydroxide (NaOH) solution (20 mL ethanol) was dropwise added under continuous stirring condition. The glass beaker was then covered tightly with aluminum foil and heated at 65°C for 2 hours with continuous stirring. After 2 h, the resultant transparent ZnO nanoparticle colloidal solution (seeding material) was allowed to cool down to room temperature and stored in a refrigerator for further use.

For seeding of the 5 nm ZnO particles on the cotton cloth, the cloths (3 cm x 3 cm) were dipped into the seeding solution for 10 mins, and then dried into a hot oven operating at 95°C for 10 mins. The procedure was repeated for 11 times. Then the cloth was transferred to autoclave vessel. An aqueous solution of zinc nitrate (20 mM) and hexamethylenetetramine (20 mM) was used as the precursor solution for the ZnO NF growth, which was carried out at 102 °C for 15 hrs. This led to the growth of ZnO NFs with a length of ~600-850 nm, and a diameter of ~150-300 nm. To maintain a constant growth rate of the ZnO NFs during the hydrothermal process, the old precursor solution was replaced with a fresh solution every 5 h. The as-obtained ZnO NF samples were then taken out of the reaction vessel and rinsed thoroughly with DI water to remove unreacted residues. Finally, the samples were annealed in air at ~110 °C for 3 h prior to the study.

### Characterization methods

Field Emission Scanning Electron Microscopy (FESEM, QUANTA FEG 250) was used to investigate the surface morphology of the samples. Transmission electron microscopy (TEM) was carried out using an FEI (Technai S-Twin) instrument with acceleration voltage of 200 kV. A drop of sample (obtained by scratching the ZnO NF decorated cotton cloth) was placed on a carbon-coated copper grid and particle sizes were determined from micrographs recorded at a high magnification of 100000X. X-ray diffraction (XRD) was used to characterize crystal phase by a PANalytical XPERTPRO diffractometer equipped with Cu Kα radiation (at 40 mA and 40 kV) at a scanning rate of 0.02° S^−1^ in the 2θ range from 20° to 80°. Thermal gravimetric analysis (TGA) of PDPB, PDPB-ZnO and ZnO solid powder was performed under nitrogen atmosphere with a heating rate of 10 °C-min from 30 °C to 600 °C by using a Perkin-Elmer TGA-50H. For optical experiments, the steady-state absorption and emission were recorded with a JASCO V-750 spectrophotometer and a JASCO FP-8200 fluorimeter, respectively. Picosecond-resolved spectroscopic studies were carried out using a commercial time correlated single photon counting (TCSPC) setup from Edinburgh Instruments (instrument response function, IRF = 82 ps, excitation at 375 nm). The details of experimental set up and methodology were described in our earlier report [72]. The average lifetime (amplitude-weighted) of a multi-exponential decay is expressed as

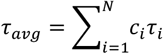

### Computational study

#### Binding study

The optimized orthorhombic ZnO wurtzite unit cell (a=5.63, b=3.25, c=5.21) with 8 atoms was taken to build a large supercell. For binding calculations on the <002>, <100>, and <101> facets, slabs of atoms with the exposed <002>, <100>, and <101> phases, respectively, were cut out of the bulk (supercell). For the calculation with a nanosphere, a sphere or radius 15 Å was cut out from the bulk and then appropriate surface atoms were removed to make it stoichiometrically neutral. ACE2 bound SARS-CoV-2 spike protein structure (PDB ID: 6lzg) was obtained from Protein Data Bank (PDB). ACE2 protein was removed before docking with the nanostructures. The structure of the lectin protein (PDB ID: 4ywa) of *P. aeruginosa* was also obtained from PDB. Molecular docking was performed using AutoDock Vina [39] following previously published protocol [40]. The docked structures were energy minimized under OPLS (optimized potential for liquid simulation) using Desmond molecular dynamics program [41] as implemented in Schrodinger Maestro (Academic release 2018-4) followed by rescoring with AutoDock Vina. Changes in the atomic positions of the protein backbone after minimization was computed in terms of root mean square deviation (RMSD).

#### Sequence analysis

Sequence of the cell surface adhesion lectin protein lecA of *Pseudomonous auregionosa* was obtained from Protein Data Bank (PDB ID: 4YWA). Structure and sequence of the receptor binding domain (RBD) of SARS-CoV-2 was also obtained from the PDB (ID: 6LZG). Sequence alignment was performed using Clustal Omega [73] with no end gap penalty. Local alignments were performed with LALIGN [74]. Similar amino acids in the binding sites of the two proteins are mapped using PyMOL molecular graphics software.

### Antibacterial tests

The bacterial strains of gram-negative *P. aeruginosa* as used in this study were obtained from Dey’s Medical Stores Mfg. Ltd., Kolkata, India. The glass wares, suction nozzles, and culture medium were sterilized in an autoclave at a high pressure of 0.1 MPa and a temperature of 120°C for 30 min before experiments. Bacteria cultures were cultivated in sterilized Luria-Bertani (LB) broth (Himedia, India) and incubation was done at 37°C with a shaking incubator for 24 hrs.

The colony count method was used to estimate antibacterial properties through the concentration of the survival colonies bacteria in co-cultured solution. First, original bacterial suspensions were washed three times with phosphate-buffered saline (PBS; pH 7.4) solution to a concentration of 10^8^ CFU/ml. Then, they were poured onto the ZnO NF functionalized cotton fabric and incubated for 1 hour. Third, the incubated bacterial solution was diluted five times to a certain concentration. The resulting bacterial PBS suspensions (100 μL) were spread on gelatinous LB agar plates, culturing at 37°C for 24 hours. The number of survival colonies was counted manually. For growth curve analysis, the five times diluted bacterial solution was added to LB and kept at 37°C. 600 nm absorbance (or, optical density; OD_600_) was monitored at regular interval (1 h) to assess the growth kinetics.

The bacteria cells after proper incubation with the ZnO NF functionalized cotton fabric were stained with DAPI and PI. The DAPI stains all cells while PI only stains the membrane disrupted cells. The 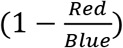 ratio was obtained to assess the survivality of *P. aeruginosa.* Non-functionalized cotton cloths and untreated bacterial cultures were used as control in all the tests. All tests were repeated at least six times.

### Statistics

Data are represented as Mean ± Standard Deviation (SD), unless otherwise stated. One-way analysis of variance (ANOVA) followed by Tukey’s *post-hoc* test was used to calculated differences between groups. *P*<0.05 was considered significant. For all statistical tests GraphPad Prism (v8.0) software was used.

