## Supplementary Information for "Nanoceutical Fabric Prevents COVID-19 Spread through Expelled Respiratory Droplets: A Combined Computational, Spectroscopic and Anti-microbial Study"

**Local alignment results between SARS-CoV-2 spike protein and lecA of *P. aeruginosa*.**

```
The best non-identical alignments are:     ls-w bits E(1) %_id  %_sim  alen
6LZG_B|Spike glycoprotein|Severe ac ( 209)   52 18.8   0.053 0.281 0.625   32
+-                                           45 16.9    0.19 0.243 0.622   37
+-                                           35 14.1    0.76 0.281 0.562   32
+-                                           30 12.7    0.98 0.833 1.000    6
+-                                           29 12.4    0.99 0.364 0.727   11
+-                                           29 12.4    0.99 0.500 0.562   16
+-                                           29 12.4    0.99 0.400 0.600   10
+-                                           26 11.6       1 0.214 0.571   28
+-                                           26 11.6       1 0.261 0.565   23
+-                                           26 11.6       1 0.286 0.571   14
+-                                           26 11.6       1 0.308 0.769   13

>>>4YWA_A|PA-I, 121 aa vs lalign-I20201121-055742-0421-60153165-p2m.bsequence library

>>6LZG_B|Spike glycoprotein|Severe acute respiratory syn  (209 aa)
 Waterman-Eggert score: 52;  18.8 bits; E(1) <  0.053
28.1% identity (62.5% similar) in 32 aa overlap (14-45:149-180)

            20        30        40
4YWA_A QVTSIIYNPGDVITIVAAGWASYGPTQKWGPQ
       .... ::. :..    . :.  : : :..: :
6LZG_B DISTEIYQAGSTPCNGVEGFNCYFPLQSYGFQ
      150       160       170       180

>--
 Waterman-Eggert score: 45;  16.9 bits; E(1) <  0.19
24.3% identity (62.2% similar) in 37 aa overlap (7-43:92-128)

         10        20        30        40
4YWA_A LANNEAGQVTSIIYNPGDVITIVAAGWASYGPTQKWG
       .: ...:....  :.  : .:  . .: : .  .: :
6LZG_B IAPGQTGKIADYNYKLPDDFTGCVIAWNSNNLDSKVG
             100       110       120

>--
 Waterman-Eggert score: 35;  14.1 bits; E(1) <  0.76
28.1% identity (56.2% similar) in 32 aa overlap (1-32:34-63)

               10        20        30
4YWA_A AWKGEVLANNEAGQVTSIIYNPGDVITIVAAG
       ::. . ..:  :    :..:: ..  :.   :
6LZG_B AWNRKRISNCVADY--SVLYNSASFSTFKCYG
            40          50         60
```

```
>--
 Waterman-Eggert score: 30;  12.7 bits; E(1) <  0.98
83.3% identity (100.0% similar) in 6 aa overlap (108-113:52-57)

       110
4YWA_A NSGSFS
       ::.:::
6LZG_B NSASFS


>--
 Waterman-Eggert score: 29;  12.4 bits; E(1) <  0.99
36.4% identity (72.7% similar) in 11 aa overlap (68-78:153-163)

        70
4YWA_A KIGNSGTIPVN
       .: ..:. : :
6LZG_B EIYQAGSTPCN
            160

>--
 Waterman-Eggert score: 29;  12.4 bits; E(1) <  0.99
50.0% identity (56.2% similar) in 16 aa overlap (35-48:176-191)

             40
4YWA_A SYG--PTQKWGPQGDR
       :::  ::.  : :  :
6LZG_B SYGFQPTNGVGYQPYR
         180       190

>--
 Waterman-Eggert score: 29;  12.4 bits; E(1) <  0.99
40.0% identity (60.0% similar) in 10 aa overlap (23-32:86-95)

             30
4YWA_A GDVITIVAAG
       :: .  .: :
6LZG_B GDEVRQIAPG
          90

>--
 Waterman-Eggert score: 26;  11.6 bits; E(1) <  1
21.4% identity (57.1% similar) in 28 aa overlap (82-108:103-130)

              90        100
4YWA_A FRWVAPNNVQG-AITLIYNDVPGTYGNN
        . .  :..  : .:.   :.. .  :.:
6LZG_B YNYKLPDDFTGCVIAWNSNNLDSKVGGN
            110       120       130

>--
 Waterman-Eggert score: 26;  11.6 bits; E(1) <  1
26.1% identity (56.5% similar) in 23 aa overlap (83-105:90-111)

             90       100
4YWA_A RWVAPNNVQGAITLIYNDVPGTY
       : .::... : :.     .:  .
6LZG_B RQIAPGQT-GKIADYNYKLPDDF
      90        100       110
```

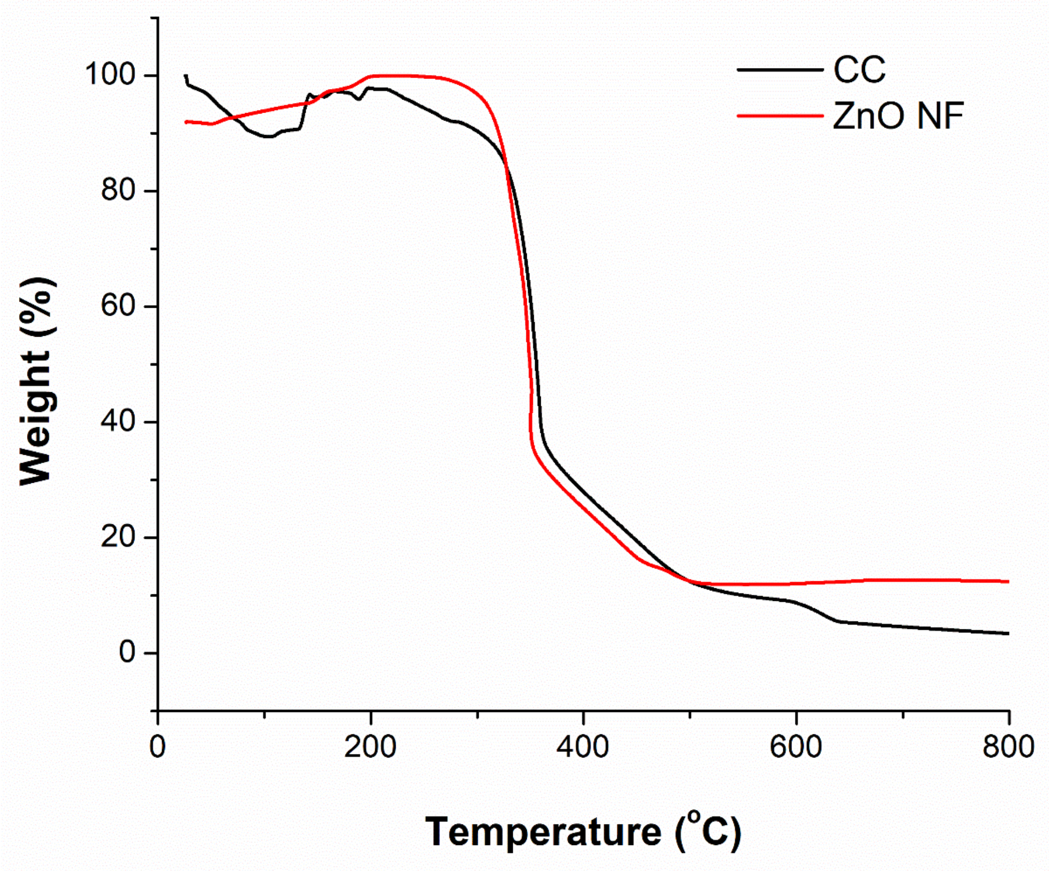


**Supplementary Figure S1. TG data of ZnO NF decorated cotton cloth and bare cotton cloth.**

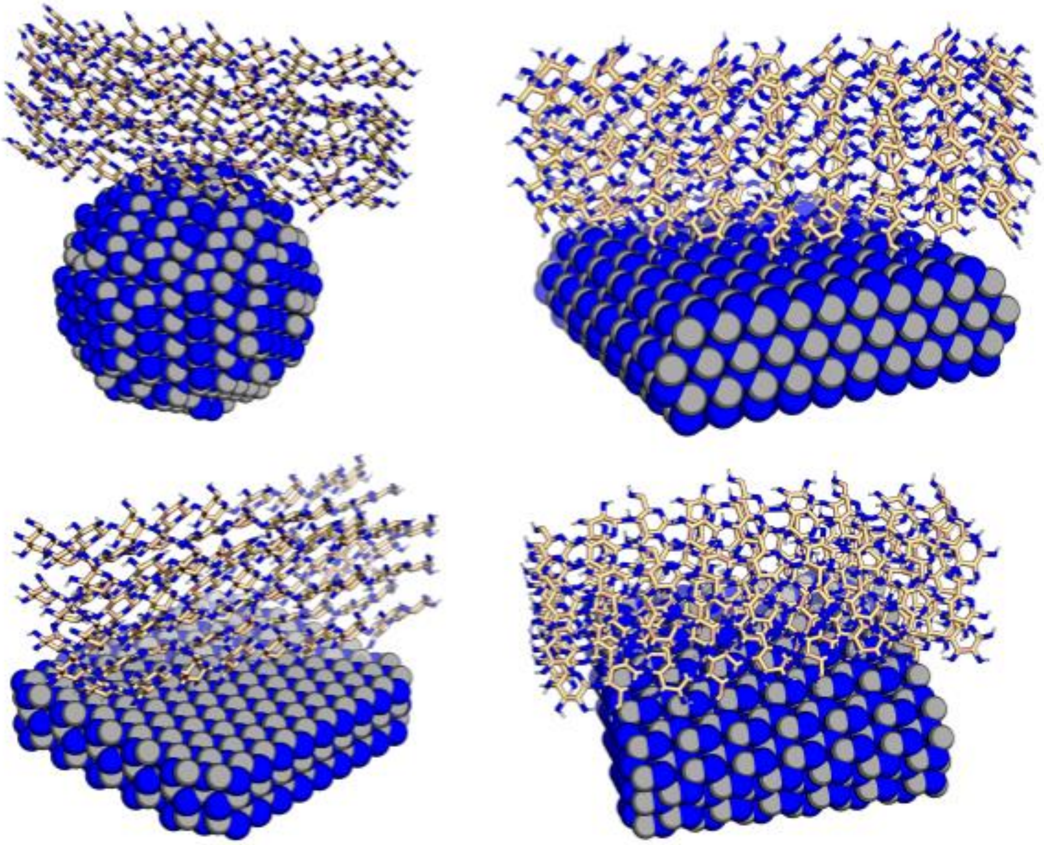

**Supplementary Figure S2. Binding of ZnO with cellulose fiber.**
